## Supplementary Information for "Functions of Direct and Indirect Pathways for Action Selection Are Quantitatively Analyzed in A Spiking Neural Network of The Basal Ganglia"

**Noname manuscript No.**
(will be inserted by the editor)

### Supplementary Information for "Functions of Direct and Indirect Pathways for Action Selection Are Quantitatively Analyzed in A Spiking Neural Network of The Basal Ganglia"

**Sang-Yoon Kim · Woochang Lim**

the date of receipt and acceptance should be inserted later

**Abstract** This is the Supplementary Information (SI) for "Functions of Direct and Indirect Pathways for Action Selection Are Quantitatively Analyzed in A Spiking Neural Network of The Basal Ganglia." In this SI, we make brief description of a spiking neural network with a single channel for the basal ganglia, considered in our prior work (Kim and Lim, 2023).

#### 1 Spiking Neural Network with A Single Channel for The Basal Ganglia

For study of action selection in the basal ganglia (BG), we consider a spiking neural network with 3 laterally interconnected channels in Fig. 1(a) (Humphries et al., 2006; Sen-Bhattacharya et al., 2018). Here, each single channel is the same as that considered in our recent works (Kim and Lim, 2023). The spiking neural network with a single channel is based on anatomical and physiological data obtained in rat-based works. Hence, we employ rat-brain terminology.

In this SI, we make brief description on the spiking neural network with a single channel. Based on the spiking neural networks for the BG developed in previous works (Humphries et al., 2009; Tomkins et al., 2014; Fountas and Shanahan, 2017), we made refinements on the spiking neural network to become satisfactory for our recent study to quantify harmony between direct and indirect pathways for the healthy and Parkinsonian states (Kim and Lim, 2023). Details on the spiking neural network with a single channel are given in Sec. II and Appendices in (Kim and Lim, 2023).

S.-Y. Kim
Institute for Computational Neuroscience and Department of Science Education, Daegu National University of Education, Daegu 42411, Korea


W. Lim (corresponding author)
Institute for Computational Neuroscience and Department of Science Education, Daegu National University of Education, Daegu 42411, Korea


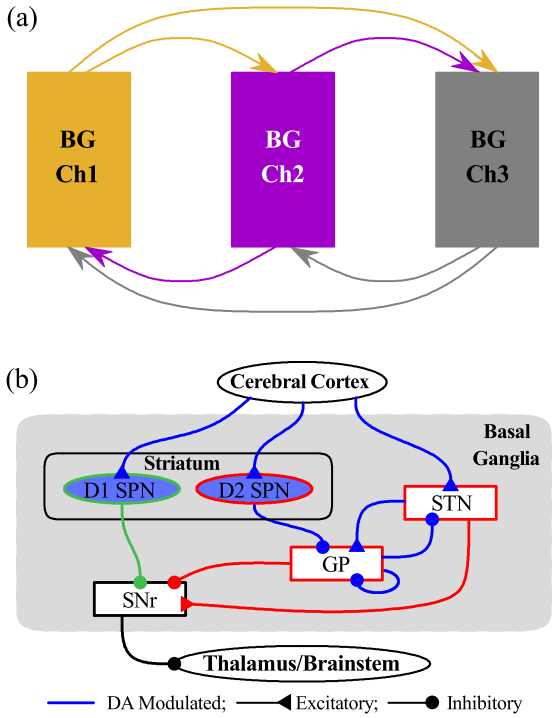


**Fig. 1** (a) Box diagram of the spiking neural network with three laterally interconnected channels for the basal ganglia (BG). The channels 1, 2, and 3 are denoted in orange, purple, and gray color, respectively. There are inter-channel connections from neighboring channels. (b) Box diagram of the spiking neural network with a single channel for the BG. Excitatory and inhibitory connections are denoted by lines with triangles and circles, respectively, and dopamine-modulated cells and connections are represented in blue color. Striatum and STN (subthalamic nucleus), receiving the excitatory cortical input, are two input nuclei to the BG. In the striatum, there are two kinds of inhibitory spine projection neurons (SPNs); SPNs with the D1 receptors (D1 SPNs) and SPNs with D2 receptors (D2 SPNs). The D1 SPNs make direct inhibitory projection to the output nuclei SNr (substantia nigra pars reticulata) through the direct pathway (DP; green color). In contrast, the D2 SPNs are connected to the SNr through the indirect pathway (IP; red color) crossing the GP (globus pallidus) and the STN. The inhibitory output from the SNr to the thalamus/brainstem is controlled through competition between the DP and IP.

Figure 1(b) shows a box diagram of major BG cells and synaptic connections in the spiking neural network with a single channel. Based on the anatomical property of the BG (Oorschot, 1996; Bar-Gad et al., 2003; Mailly et al., 2003; Sadek et al., 2007), we take into consideration the spiking neural network, consisting of D1/D2 spine projection neurons (SPNs), subthalamic nucleus (STN) neurons, globus pallidus (GP) neurons, and substantia nigra pars reticulata (SNr) neurons. For more details on the numbers of the BG cells and their synaptic connection probabilities, refer to Sec. IIA and Tables I and II in (Kim and Lim, 2023).

Next, we make brief description of the single neuron models and the dopamine (DA) effects in the spiking neural network with a single channel; for details refer to Sec. IIB and Appendix A in (Kim and Lim, 2023). As the single neuron model, we employ the Izhikevich spiking neuron model (which is not only biologically plausible, but also computationally efficient) as the elements of the spiking neural network (Izhikevich, 2003, 2004, 2007a,b). The spiking neural network consists of 5 populations of D1/D2 SPNs, STN, GP, and SNr; for parameter values of each BG cells, refer to Table III in (Kim and Lim, 2023). The modulation effect of DA on the D1/D2 SPNs are also taken into consideration (Humphries et al., 2009; Tomkins et al., 2014; Fountas and Shanahan, 2017). For details, refer to Sec. IIB, Appendix A, and Table IV in (Kim and Lim, 2023).

The state of a neuron in each population is characterized by its membrane potential and slow recovery variable. Time-evolution of the membrane potential and the slow recovery variable is governed by 3 kinds of currents into the neuron such as the external current from the external background region, the synaptic current, and the injected stimulation current. Here, we consider the case of no injected stimulation current. The external current is modeled in terms of spontaneous (in-vivo) current (to get the spontaneous in-vivo firing rate) and random background input; for more details, refer to Sec. IIB, Appendix A, and Table V in (Kim and Lim, 2023).

We also consider the synaptic currents and the DA effects in the case of single channel; detailed explanations are given in Sec. IIB and Appendix B in (Kim and Lim, 2023). There are 3 kinds of synaptic currents from a presynaptic source population to a postsynaptic neuron in the target population; 2 kinds of excitatory AMPA and NMDA receptor-mediated synaptic currents and one type of inhibitory GABA receptor-mediated synaptic current. For each $R$ (AMPA, NMDA, and GABA) receptor-mediated synaptic current, the synaptic conductance is given by a product of the maximum synaptic conductance, the average number of afferent synapses, and the fraction of open postsynaptic ion channels. The time course of fraction of open ion channels is provided by a sum of exponential functions over presynaptic spikes. The synaptic parameters are given in Table VI in (Kim and Lim, 2023). These synaptic parameter values are based on physiological property (Park et al., 1982; Nakanishi et al., 1990; Fujimoto and Kita, 1993; Góngora-Alfaro et al., 1997; Götz et al., 1997; Richards et al., 1997; Bevan and Wilson, 1999; Bevan et al., 2000; Dayan and Abbott, 2001; Bevan et al., 2002; Liu et al., 2022; Hallworth et al., 2003; Baufreton et al., 2005; Wolf et al., 2005; Shen and Johnson, 2006; Moyer et al., 2007; Gertler et al., 2008; Bugaysen et al., 2010; Connelly et al., 2010; Ammari et al., 2011). The modulation effect of DA on afferent synapses into the D1/D2 SPNs, the STN, and the GP is also taken into consideration (Humphries et al., 2009; Tomkins et al., 2014; Fountas and Shanahan, 2017); for details, refer to Table VII in (Kim and Lim, 2023).

Finally, we note that, in the spiking neural network with 3 channels, inter-channel connections are made via widespread diffusive excitation from the STN in a channel to the target nuclei, SNr and GP, in all the 3 channels (Parent and Hazrati, 1993, 1995a,b). Thus, to the target neurons, SNr and GP, in a channel, there are one intra-channel synaptic current from the source STN in the same self-channel and two inter-channel synaptic currents from the source STN in the two neighboring channels. The intra-channel synaptic current is just the same as that considered in the above case of single channel. But, unlike the case of single channel, two additional inter-channel synaptic currents from STN in the neighboring channels to the target nuclei, SNr and GP, in a channel must be considered.

The multi-channel (MCh) connection probability $p^{(T,\mathrm{STN})}_{(c,MCh)}$ from the source STN to the target (SNr, GP) in both cases of intra- and inter-channel interactions is given by $p^{(T,\mathrm{STN})}_{(c,MCh)} = p^{(T,\mathrm{STN})}_{c}/N_C$; $p^{(T,\mathrm{STN})}_{c}$ is the connection probability in the case of single channel and $N_C$ is the number of channels (Humphries et al., 2006). Then, the number of afferent synapses into the target neurons becomes constant, independently of $N_C$. In the present work, $N_C = 3$.
